# Necklace: combining reference and assembled transcriptomes for more comprehensive RNA-Seq analysis

**DOI:** 10.1101/200287

**Authors:** Nadia M Davidson, Alicia Oshlack

## Abstract

**Background:** RNA-Seq analyses can benefit from performing a genome-guided and de novo assembly, in particular for species where the reference genome or the annotation is incomplete. However, tools for integrating assembled transcriptome with reference annotation are lacking.

**Findings:** Necklace is a software pipeline that runs genome-guided and de novo assembly and combines the resulting transcriptomes with reference genome annotations. Necklace constructs a compact but comprehensive superTranscriptome out of the assembled and reference data. Reads are subsequently aligned and counted in preparation for differential expression testing.

**Conclusions:** Necklace allows a comprehensive transcriptome to be built from a combination of assembled and annotated transcripts which results in a more comprehensive transcriptome for the majority of organisms. In addition RNA-seq data is mapped back to this newly created superTranscript reference to enable differential expression testing with standard methods. Necklace is available from https://github.com/Oshlack/necklace/wiki under GPL 3.0.

## Findings Introduction

Despite the increasing number of species with a sequenced genome, the vast majority of reference genomes are incomplete. They may contain gaps, have unplaced assembly scaffolds and be poorly annotated. Therefore, analyzing RNA-Seq data using just the reference genome has the potential to miss important biology for many organisms. Ideally, an RNA-Seq analysis could utilise prior knowledge of the gene-models available from a reference genome and annotation, whilst also extracting information about the genes from the data itself through genome-guided and/or de novo assembly [1,2].

In [3] we introduced the concept of the superTranscriptome, where each gene is represented by one sequence containing all of that gene’s exons in transcriptional order. SuperTranscripts provided a convenient means in which transcriptomes from difference sources, such as assembly and annotation, can be combined into a compact and unified reference. When applied to chicken, we showed that we could recover hundreds of segments of genes that were absent from the chicken reference genome.

Here we present software called necklace which automates the process described in [3] for any species with an incomplete reference genome. Necklace takes as input a configuration file containing paths to the RNA-Seq reads, a reference genome and one or more reference genome annotation. Because de novo assembly is error prone, we require that any gene discovered specifically through de novo assembly be also found amongst the coding sequence of a related (well annotated) species. Therefore, the genome and annotation of a related species must also be provided to necklace. Necklace will then run the steps involved in genome-guided and de novo assembly, and combine the assembled transcriptome with reference annotations for the species of interest After building the superTranscriptome, necklace will align and count reads in preparation for testing for differential gene expression and differential transcript usage using well established tools such as edgeR [4], DEseq[5] or DEXseq[6].

In order to demonstrate the application of necklace in a new data set we analysed public RNA-seq data from sheep milk. Compared to using the sheep reference genome on it own, the necklace analysis resulted in 18% more reads being assigned to genes and 19% more differentially expressed genes being detected.

### The Necklace pipeline

Necklace is a pipeline constructed using the bpipe framework [7]. It steers external software, such as aligners and assemblers, as well as a set of its own utilities, written in c/c++. As input Necklace takes the raw RNA-seq reads and the reference genome for the species as well as any available annotation. In addition, it takes a reference genome and annotation from a related, but well studied species such as human, drosophila, yeast, etc. Necklace consists of several sequential stages: initial genome guided and *de novo* assembly, clustering transcripts into gene groupings, reassembly to build the superTranscriptome and finally alignment and counting of mapped reads in preparation for differential expression testing and differential isoform usage testing. Each of these sequential stages consists of several sub-stages and is outlined in Figure 1 with further detail below.

1. **Assembly** – The assembly stage creates three different transcriptomes. First reads are aligned to the reference genome using HISAT2 [8] and genome-guided assembly is performed with StringTie [9]. This assembly is combined with reference annotations and then flattened based on genomic location, so that each exon is reported only once and overlapping exons are merged. Exonic sequence is then extracted from the genome and concatenated to build a “genome-based” superTranscriptome. In parallel, the related species annotation is used to create a “genome-based” superTranscriptome (without genome-guided assembly). Finally, RNA-Seq reads are de novo assembled with Trinity [10].
2. **Clustering of transcripts** – This step assigns de novo assembled transcripts to gene clusters prior to building the final superTranscriptome. Those contigs aligning to the genome-based superTrascriptome (using Blat [11]) are allocated to known genes while those not aligning to the genome, but found in the related species superTranscriptome are assigned to novel genes. De novo assembled transcripts that align to more than one gene are removed to avoid false chimeras [12] from being introduced into the superTranscriptome.
3. **Reassembly of superTranscripts** – Each cluster consists of a gene’s genome-based superTranscript and/or its set of de novo assembled transcripts. The transcripts in each cluster are merged together through Lace assembly [3], to produce one superTranscript per gene.
4. **Summarization** – Reads are aligned back to the superTranscriptome using HISAT2 and fragments counted per gene using featureCounts [13]. Splice junctions reported by HISAT2 are used to segment each superTranscript into a set of contiguous “blocks”. Fragments are then counted in “blocks” and can be used for differential isoform detection like exon counts.

**Figure 1.**
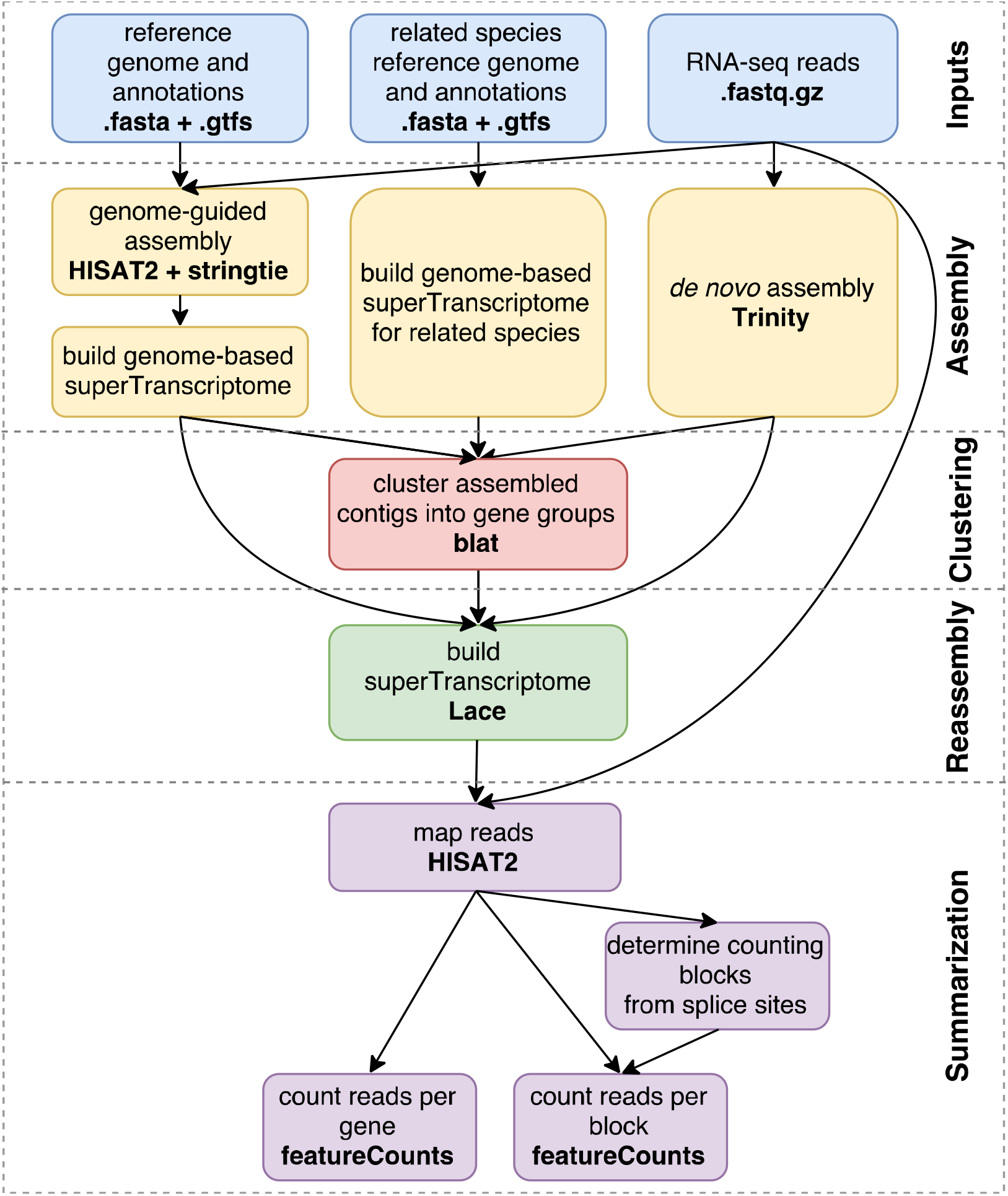
An overview of the necklace pipeline. External software that necklace runs is shown in bold.

### Application to differential expression testing in sheep transcriptomes

To demonstrate the utility of necklace, we applied it to public RNA-Seq from Churra sheep milk and compared transcriptome expression at day 10 to day 150 post lambing [14], Necklace was given the sheep reference genome, Oar_v3.1. Human, with the hg38 reference genome, was used as the related species. For both genomes, version 90 of the Ensembl annotation was used (see methods). Using this data and set of reference files resulted in a more comprehensive transcriptome. Compared to the Ensembl sheep annotation, the number of bases included in the necklace transcriptome increased by 76% and there were 2208 (8%) more genes identified. This more comprehensive reference resulted in 18% more reads being assigned to genes by featureCounts compared to the reference annotation alone. When performing differential expression analysis using edgeR the necklace transcriptome identified more significantly differentially expressed genes (456 compared to 383, FDR<0.05). Some of these differences could be attributed to the inclusion of novel unannotated genes with 66 of the newly annotated genes being identified as differentially expressed. Necklace was also able to improve the detection of differential expression amongst several known genes by providing more complete gene sequences. Larger numbers of reads mapping to the longer sequences resulted in more power for differential expression testing. One example of this was the *SERTM1* gene where the annotated transcript only included 321bp while the necklace superTranscript contained 3333bp and overlapped a genome assembly gap [Figure 2).

**Figure 2.**
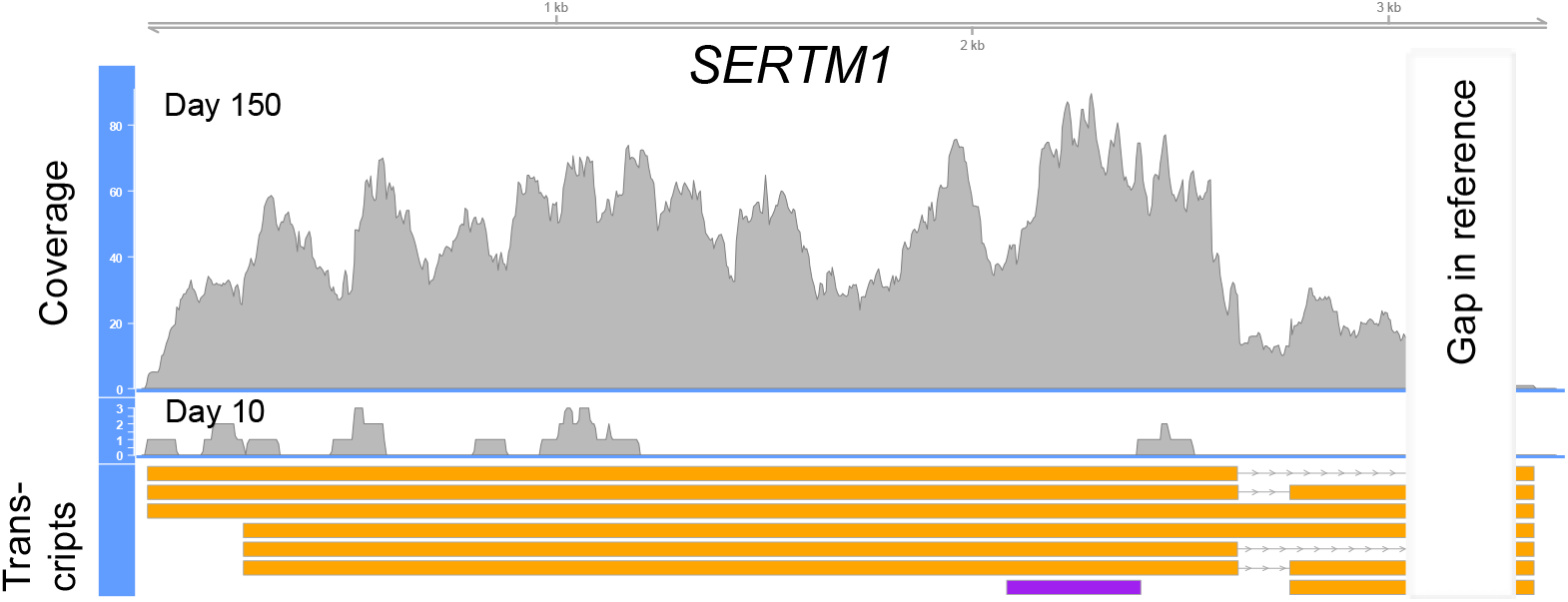
Read coverage aggregated over replicate samples for the necklace assembled superTranscript of SERTM1. This gene is found to be significantly differentially expressed using the necklace generated reference, but is missed when the reference genome and annotation are used in isolation due to low read counts. The reference annotation consists of a single transcript of 321bp (shown in purple), whereas the de novo assembled gene consists of seven transcripts up to 3333bp long (shown in orange) and includes approximately 250bp that is absent from the reference genome, in a location consistent with a genome assembly gap. The genome-guided transcripts that were assembled for this gene were filtered out by stringTie’s merge function due to an average FPKM < 1.

## Conclusion

Here we have presented necklace, a pipeline designed to improve RNA-Seq analysis in species with an incomplete genome and annotation. We believe necklace is the first pipeline to automate the steps required to combine reference and assembled data, alignment and summarization of counts. We show that this process results in more complete transcriptomes using a sheep data set and superTranscripts give more power for differential expression analysis.

## Methods

Sheep RNA-Seq data was downloaded from SRA (accession numbers SRR2932539-SRR2932542,SRR2932561-SRR2932564). The sheep genome and annotation was downloaded from Ensembl: ftp://ftp.ensembl.org/pub/release-90/fasta/ovisaries/dna/Ovisaries.Oarv3.1.dna.toplevel.fa.gz ftp://ftp.ensembl.org/pub/release-90/gtf/ovis_aries/Ovis_aries.Oar_v3.1.90.gtf.gz

The human reference genome and annotation was also downloaded from Ensembl: ftp://ftp.ensembl.org/pub/release-90/fasta/homo_sapiens/dna/Homo_sapiens.GRCh38.dna.toplevel.fa.gz. ftp://ftp.ensembl.org/pub/release-90/gtf/homo_sapiens/Homo_sapiens.GRCh38.90.gtf.gz

We then selected coding sequence from the human annotation using the command:

~~~
*grep “CDS” data/Homo_sapiens.GRCh38.90.gtf >
Homo_sapiens.GRCh38.90. CDS.gtf*
~~~

For the necklace analysis of sheep milk, all data files were placed into a subdirectory called “data” and a necklace input file, “data.txt”, was created with the following lines:

~~~
*//sequencing data
reads_R1=“data/SRR2932539_1.fastq.gz,data/SRR2932540_ ī,fastq.gz,data/SRR29
3254 1._ 1.fastq.gz,data/SRR2932542_1.fastq.gz,data/SRR2932561_ 1.fastq.gz, data/
SRR2932562_1.fastq.gz,data/SRR2932563_1.fastq.gz,data/SRR2932564_1.fastq.gz
"
reads_R2=“data/SRR2932539_2.fastq.gz,data/SRR2932540_2.fastq.gz,data/SRR29
3254 1._2.fastq.gz,data/SRR2932542_2.fastq.gz,data/SRR2932561_2.fastq.gz,data/
SRR2932562_2.fastq.gz,data/SRR2932563_2.fastq.gz,data/SRR2932564_2.fastq.gz
"

//The genome and annotation
annotation=“data/Ovis_aries.Oar_v3.1.90.gtf”
genome=“data/Ovis_aries.Oar_v3.1.dna.toplevel.fa”

//The genome and annotation of a related species
annotation_related_species=“data/Homo_sapiens.GRCh38.90.CDS.gtf”
genome_related_species=“data/Homo_sapiens.GRCh38.dna.toplevel.fa”*
~~~

Necklace version 0.9 was then run using the command:

~~~
*<necklace path>/tools/bin/bpipe run-n 8 <necklace path>/necklace.groovy
data/data.txt*
~~~

Version numbers of all the external tools that necklace calls can be found in necklace’s installation script, “install_linux64.sh”.

To make the reference based analysis as similar as possible to the necklace pipeline we used the versions of HISAT2, samtools and featureCounts that were installed by necklace.

HISAT2 was run on each sample using the command:*hisat2 --known-splicesite-infile <splice sites file>-x <genome index>-1 <input_1.fastq.gz> -2 <input_2.fastq.gz> | samtools view -u -> <output.bam>*

Where the splice sites file and genome index were the same ones generated in the initial stage of necklace that aligns reads to the reference genome.

Reads were then counted for each annotated gene using featureCounts with the command:

~~~
*featureCounts-T 8 --primary -p -t exon -g gene_id -a Ovis_aries.Oar_v3.1.90.flat.gtf -
0 counts *.bam*
~~~

Where “Ovis_aries.Oar_v3.1.90.flatgtf” was a flattened version of the sheep Ensembl annotation and was created with the necklace command:

~~~
*gtβflatgtf Ovis_aries. Oar_v3.1.90.gtf Ovis_aries. Oar_v3.1.90.flat.gtf*
~~~

Flattening the annotation involves merging transcripts of a gene into a non-redundant but complete set of exons.

For differential gene expression testing, gene-level counts were analysed using the R bioconductor package edgeR [version 3.18.1] [15]. We modeled both the time-point post lambing and animal in the design matrix:

~~~
*time_point<-c(rep (“Dayl 0”, 4), rep (“Day150”, 4))
indv<-c(3141,4860,49537,9539,3141,4860,9539,49537) //numbers are animal IDs
design <-model.matrix(~0 +factor(indv) +factorftim e_point))
colnames(design) <-gsub(“factor”, “”,colnames(design))
sample_ names=paste(indv, time_point,sep=“_”)
rownames(design)=sample_names*
~~~

The counts table was read into R and passed to edgeR:

~~~
*counts=count_table[,7:ncol(count_table)]
y <-DGEList(counts=counts)*
~~~

Genes with a counts per million [cpm] less than or equal to 0.5 in less 4 samples were filtered out and the libraries normalized.

~~~
*keep <-rowSums(cpm(y) > 0.5) >=4
y <-y[keep,, keep.lib.sizes=TRUE]
y <-calcNormFactors(y)*
~~~

We then estimated the dispersion and looked for differential expression with a false discovery rate [FDR] < 0.05:

~~~
*y <-estimateDisp(y,design,robust=TRUE)
fit <-glmFit(y, design,robust=TRUE)
qlf<-glmLRT(fit,coef=5)
is.de <-decideTests(qlf, p.value=0.05)*
~~~

## Availability of supporting source code and requirements

Project name: Necklace

Project home page: https://github.com/Oshlack/necklace/wiki

Operating system(sĵ: Linux

Programming language: Groovy and C/C++

Other requirements: Java 1.8 License: GPL 3.0

## Declarations

### List of Abbreviations

FPKM: fragements per kilobase of exon per million mapped reads

## Competing interests

None declared.

## Funding

AO is funded by an *NHMRC CDF GNT1126157*.

## Authors’ contributions

ND wrote all the software and drafted the paper. AO oversaw the project and contributed to writing the manuscript.

## Acknowledgements

We would like to thank Anthony Hawkins, the author of Lace, who contributed to the early concept of necklace when applied to chicken.

